# A microfluidic module for real-time generation of complex multi-molecule temporal concentration profiles

**DOI:** 10.1101/119701

**Authors:** Kristina Woodruff, Sebastian J. Maerkl

## Abstract

We designed a microfluidic module that generates complex, dynamic concentration profiles of multiple molecules over a large concentration range using pulse-width modulation (PWM). Our PWM device can arbitrarily combine up to 6 different inputs and select between three downstream mixing channels as required by the application. The module can produce arbitrary concentrations with a dynamic range of up to 3-5 decades. We created complex concentration profiles of 2 molecules, with each concentration independently controllable, and show that the PWM module can execute rapid concentration changes as well as long-timescale pharmacokinetic profiles. Concentration profiles were generated for molecules with molecular weights ranging from 560 Da to 150 kDa. Our PWM module produces robust and precise concentration profiles under a variety of operating conditions, making it ideal for integration with existing microfluidic devices for advanced cell and pharmacokinetic studies.

## Introduction

In order to perform complex and biologically relevant experiments on microfluidic platforms, there is a need for accurate and automated methods to manipulate molecular concentrations on chip. Microfluidic devices are generally connected to a small number of input solutions. Consequently, most experiments involve one or more step function changes, switching rapidly from one molecule to another or from one concentration to another. In contrast, naturally occurring changes in concentrations of molecules are rarely instantaneous step-functions. Physiologically relevant changes, such as pharmacokinetic drug concentration profiles, occur over minutes to hours and follow a complex, continuous rise and fall pattern. Complex temporal concentration profiles of one or more substances could be used to study the influence of changing antibiotic concentrations on bacteria, the effects of drugs on mammalian cells, and stem cell differentiation. The ability to rapidly generate arbitrary and complex temporal concentration profiles is thus of general interest and a useful experimental method.

One technique to create concentration profiles from two inputs involves lateral diffusion-based gradients^1,2^. Two fluids are simultaneously flowed parallel to one another along the length of a channel or chamber, establishing a gradient that is perpendicular to the direction of the flow. This principle has been used to design microfluidic devices capable of preparing up to 81 chemical combinations^3^. However, the total possible outputs are highly dependent on the number of inputs (16 stock solutions are needed to produce 81 different solutions) and because this approach is diffusion-based, it offers poor spatio-temporal resolution. Several active mixing techniques^4^ including mechanical micromixers^5^, microstructures^6^, integrated peristaltic pumps^7^, and serial dilution schemes^8^ have been developed to provide fast concentration changes. As with the gradient-based technique, the number of possible concentrations on these devices is also limited and defined by the number of solution inputs.

To generate small changes in concentration over a large dynamic range, dynamic strategies such as pulse-width modulation (PWM) are necessary. PWM is based on the concept of controlling the duty cycle of a load to control voltage and current in electrical engineering. The microfluidic equivalent controls the flow of buffer and substrate reservoirs. A microfluidic device that supports fast switching times can generate alternating pulses of buffer and substrate. When directed through a channel, these pulses diffuse to homogeneity. Total cycle time (time to execute one pulse of buffer and one pulse of substrate) is kept constant. Different concentrations are created by varying the duty cycle, which refers to the fraction of time occupied by the substrate pulse in comparison to the total cycle time.

PWM has been incorporated into microfluidic diluters in the past^9,10,11^. For a single molecule of small size, previous examples have prepared solutions of specific concentrations with high accuracy. The compatibility of these devices with proteins and substances larger than small dye molecules has not been tested. Moreover, current PWM devices have not demonstrated that complex patterns can be generated for multiple molecules simultaneously. Another drawback is the large, non-standard dimensions of the mixing infrastructure; these elements require additional cleanroom fabrication steps and occupy a significant amount of space on the chip. Lastly, previously reported PWM chips have performed experiments lasting from a few minutes to 1.5 hours. It is unknown whether these devices are capable of sustaining concentrations for extended time periods, as experiments spanning several hours or days would be required for studies involving bacteria or mammalian cells.

Long-term concentration manipulation on microfluidic devices would also be interesting for pharmacokinetic and pharmacodynamic (PK/PD) studies. These assays are central to drug discovery and development ^12,13^. Typical *in vitro* PK/PD models, while able to properly reproduce the pharmacokinetics of drugs *in vivo*, provide only bulk measurements ^14^. This simplified setup poorly represents the complex *in vivo* environment and is not compatible with techniques that probe single cell phenotypes. Although microfluidic chips have been designed to address this need^15,16^, these devices lack the ability to simulate the gradually rising and falling concentrations of drugs in plasma. Ideally, a device should be able to create realistic PK/PD profiles for multiple substances at once.

We have developed a microfluidic device that improves upon previous PWM-based chips, making it compatible with a broad range of applications. Our device successfully implements concentration changes over both short and long time scales, making it capable of producing complex PK/PD profiles. The device occupies minimal space and its features can be fabricated in one photolithographic layer. These characteristics facilitate integration of the module into existing chip designs. We also show that the PWM device can function as an independent module connected upstream of a second chip. The ability to easily modify total cycle times and choose between 3 mixing channel options (short serpentine, long serpentine, and tubing connection to a second chip) ensure that our chip can function with a variety of molecule sizes and flow rates. Two substances can be manipulated in parallel, and the chip supports a dynamic range of nearly 3 orders of magnitude in output concentrations. These qualities make our chip an attractive tool for chemical and biological research requiring complex profiles and will enable studies that were previously inaccessible because of technical limitations.

## Results

### PWM chip design and technique

We designed a microfluidic module (Fig. 1a) that performs PWM to mix up to 6 different input solutions. The device measures 1.2 × 2.8 cm and was fabricated using standard photolithography and soft lithography techniques. The valves controlling the flow of these inputs were engineered for fast actuation (<30 ms, Supplementary Fig. 1) in order to accurately dispense short pulses of each input solution. The input channels merge into a common channel, and the flow can be directed into one of three paths depending on the application. A short serpentine channel (53 mm length, 0.074 μL capacity) facilitates diffusion-based mixing of small molecules and most proteins as they travel along the channel. The long serpentine channel (255 mm length, 0.356 μL capacity) provides for the mixing of complex solutions or large proteins with small diffusion coefficients. Lastly, flow can be directed into a short channel (8 mm length) and subsequently connected to another chip via flexible tubing. Merging of the pulses will occur in the tubing, delivering a homogeneous solution to the connected chip.

**Figure 1.**
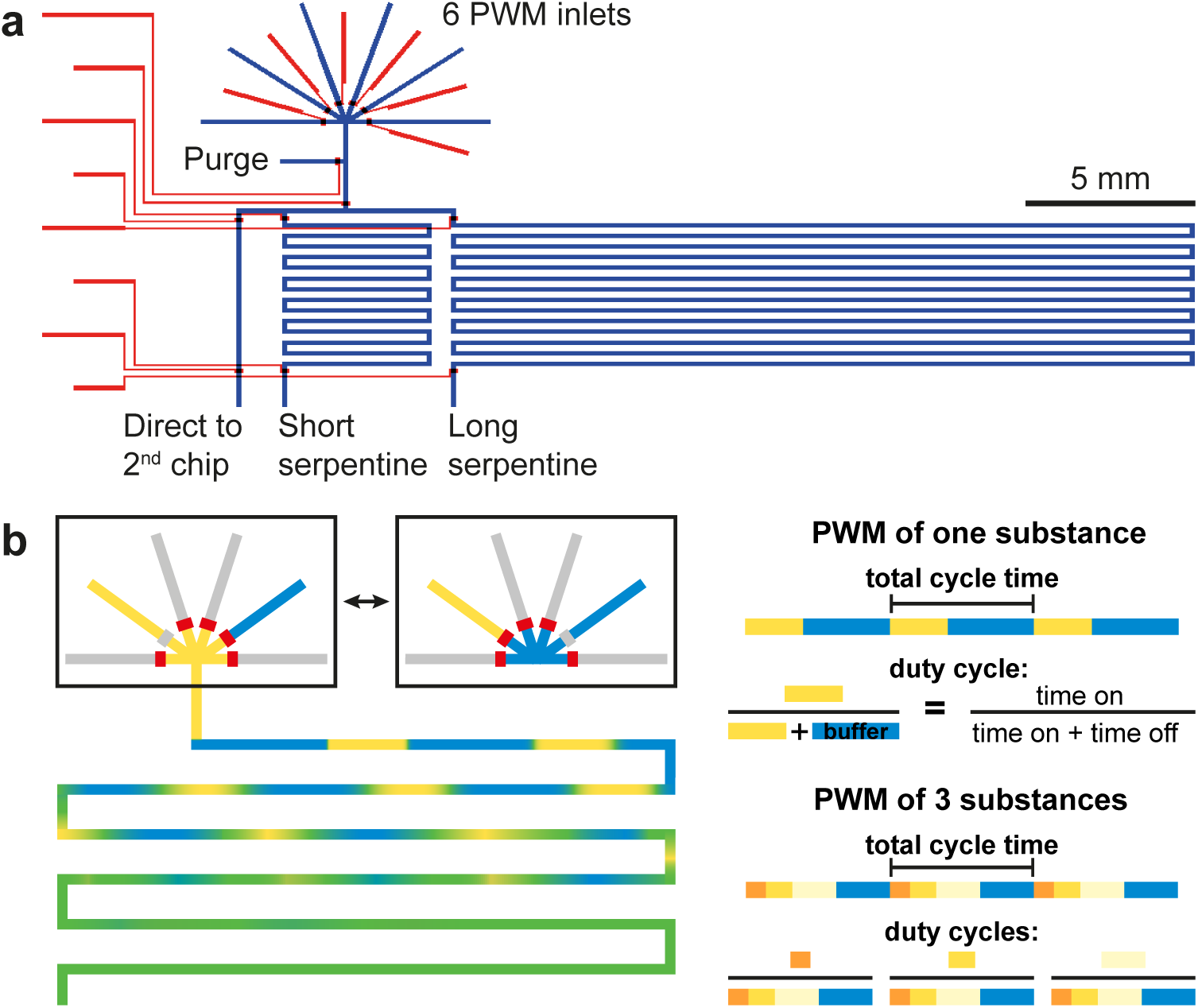
The PWM platform. (a) Design of the PWM mixing chip. Red, control layer for valve actuation; blue, flow layer for fluid manipulation. (b) Schematic of the PWM technique. Pulses are created by alternating the opening and closing of inlets. The pulsed flow pattern diffuses to homogeneity in the serpentine channels.

To adjust the concentration generated on chip, we modified the amount of time that each solution was flowed to produce pulses of different lengths. Only one solution was flowed at a time, and both the flow rate and the total cycle time (time to complete one round of pulses) were kept constant (Fig. 1b). Of the 6 inputs, at least one was a buffer used for dilution. The other 5 can be either 5 different substances (to prepare a complex mixture), the same substance diluted to 5 different concentrations (to enable an expansive dynamic range), or any combination of these two cases. We wrote a program that selects which inputs to use and executes valve opening and closing to create pulse patterns that reflect the programmed profile.

### Optimizing PWM total cycle length

To determine the optimal operating conditions for the PWM chip, we conducted experiments with 2 inputs. Since we aimed to engineer a chip that would be functional for low and high molecular weight proteins in addition to small molecules, we tested 10 kDa FITC-dextran, 150 kDa FITC-dextran, and 559 Da sulforhodamine as substrates (Fig. 2a). 9 different duty cycles were consecutively implemented on chip using total cycle times of 0.5, 1, 2, and 4 s (Fig. 2a). Both the short and long serpentine channels were tested, with flow rates of 24 μL/h and 19 μL/h, respectively. The fluorescence intensity of the fluid was measured at the end of the mixing channels and analyzed.

**Figure 2.**
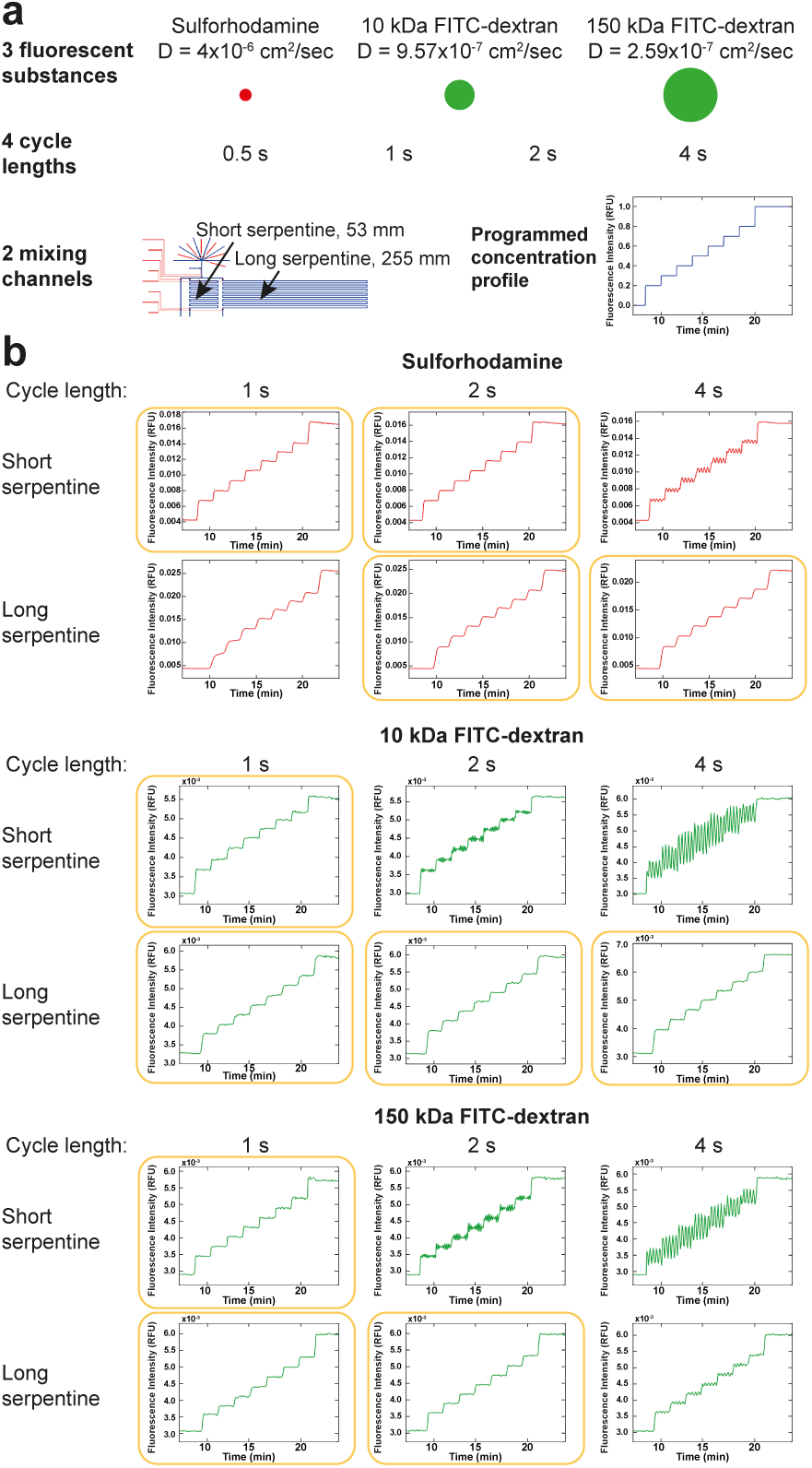
Determining optimal PWM conditions. (a) Schematic of experimental conditions tested and programmed concentration profile. (b) Experimental results. Images were acquired every 5 s at the end of the serpentine channels and analyzed for fluorescence intensity. Rightward shift of the data compared to the expected profile is caused by the time required for the solutions to travel from the PWM area to the end of the channel. Yellow boxes indicate optimal conditions.

We observed repeated and random experimental failures when using a 0.5 s cycling time (Supplementary Fig. 2). This result could possibly arise from the inability of the automated setup (a LabVIEW-controlled relay board and solenoid valves) to perform accurately over extended periods of time with such high switching rates. Alternatively, the 30 ms valve opening/closing time may be non-negligible in comparison to the short pulse lengths. All other cycle times (1-4 s) yielded an excellent linear correlation between programmed duty cycle and mean fluorescence intensity for both the short and long serpentine and for all 3 fluorescent substances (Fig. 2b). The concentrations produced on chip (Fig. 2b) closely match the programmed profile (Fig. 2a).

In addition to establishing that concentration can be precisely regulated, we tested conditions that gave rise to the most homogeneous solutions. In cases of insufficient diffusion, the pulses generated at the inlet are not completely mixed by the time they reach the measurement area. These pulses manifest as spikes of alternating high and low intensity on the raw data graphs (Fig. 2b) and equate to sizeable standard deviations on the extrapolated duty cycle graphs (Supplementary Fig. 3). Mixing is presumably more restricted for larger substrates, shorter mixing channels, or longer pulses created by longer cycle times. We observed these trends in our data; distinct pulses were especially perceptible for the 10 and 150 kDa dextrans flowed through the short serpentine at increased (2-4 s) cycle lengths (Fig. 2b). By testing this matrix of various operating conditions, we identified the combinations of molecule size, mixing channel, and cycle length for optimal performance.

### Characterizing response time

We next conducted experiments to determine the response time of our PWM module, defined as the amount of time needed to switch from 10 to 90% of the maximum concentration. We used a flow rate of 36 μL/h for the short mixing channel and 30 μL/h for the long channel. When performing PWM with a 1 s total cycle time, we obtained response times as low as 5 s (Fig. 3a,b). Depending on which substance was used, response times for the long mixing channel were 2.3-3x greater than those of the short channel. Descending from 90 to 10% concentration required slightly more time than rising from 10 to 90%. As anticipated, response time increased as substrate molecular weight decreased; for large diffusion coefficients, steep ramps became spread out due to extensive diffusion (Fig. 3a). Accordingly, the short serpentine is better suited for experiments that call for sudden and extreme changes of diffusion-sensitive small molecules. However as shown in Fig. 2b and Supplementary Fig.3, the longer serpentine can be successfully used for larger molecules and to create profiles that entail more gradual changes.

**Figure 3.**
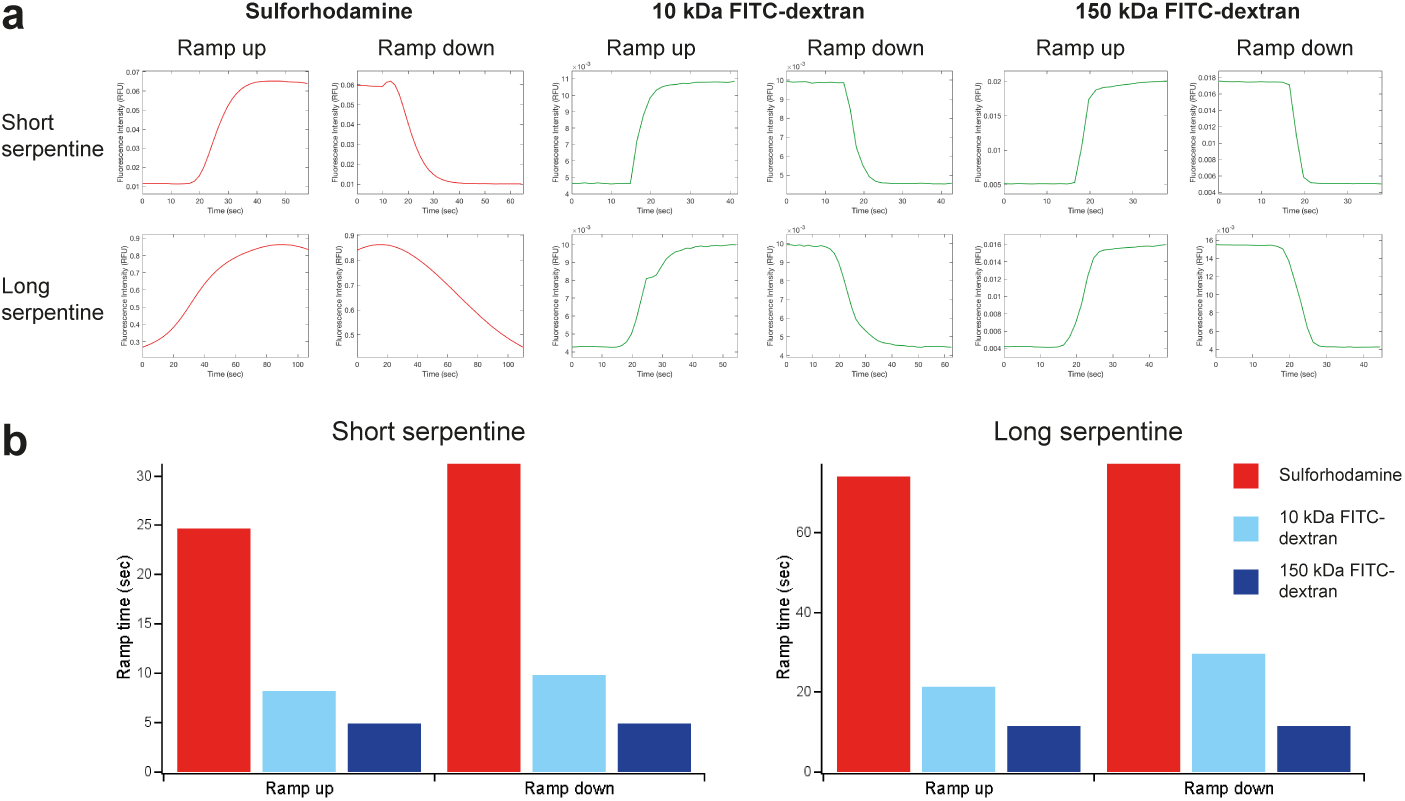
Response times required to switch between 10 and 90% of the maximum concentration. (a) Plots showing the time needed to ramp up and down. (b) Summary of results.

### Testing different flow rates on the PWM module

The data presented in Figures 2, 3, and Supplementary Figure 3 attests to the precision and quick response time of our chip when operated with typical flow rates. For these experiments, we used flow rates of ~24-36 μL/h for the short channel and ~19-30 μL/h for the long channel, generated by pressurizing the flow inputs with 4 or 10 psi, respectively. These pressures are within the standard range used for microfluidic chips of similar dimensions ^17,18^. To ensure the compatibility of our chip with a greater variety of applications, we also tested the chip with faster and slower flow velocities.

Slow flow rates (11-16 μL/h) worked best when the short mixing channel was used with a total cycle length of 1-2 s (Fig. 4, Supplementary Fig. 4). Using a long mixing channel with a slow flow rate bears the risk that the fluid will spend too much time in the channel. Subsequently, extensive diffusion will obscure the programmed concentration steps. We mitigated this effect in the long channel by extending the total cycle time to 4 s in order to create large pulses that would homogenize more slowly. This strategy would work best for small (~10 kDa) proteins; the 150 kDa FITC-dextran pulses did not completely combine, and the sulforhodamine concentration steps were blended as a consequence of diffusion (Fig. 4, Supplementary Fig. 4).

**Figure 4.**
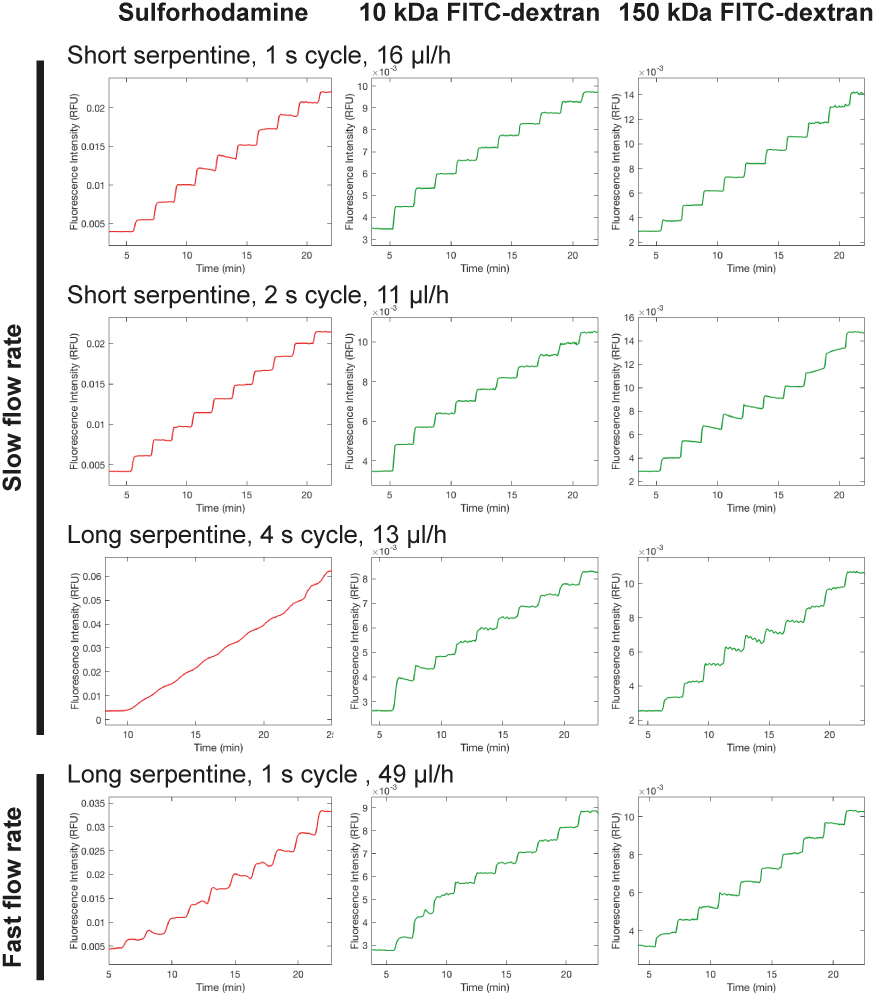
Measured fluorescence intensity resulting from step-wise duty cycle increases over time. Images were acquired at the end of the serpentine channels every 5 s and analyzed for fluorescence intensity.

When coupling faster flow rates (49 μL/h) with the short mixing channel, pulses passed through the channel too quickly and exited still intact. Only the long channel promoted complete diffusion for fast flow rates owing to the increased passage time (Fig.4, Supplementary Fig. 4). Additionally, we applied the shortest possible total cycle time (1 s) to generate short pulses that would homogenize faster and expedite the mixing process. Comparing the fast flow data to the slow flow data reveals that our chip can support more than a 4-fold change in flow rates. These flow rates (11-49 μL/h) are compatible with those used for low to medium throughput microfluidic studies of bacteria ^19,20^ and mammalian cells ^21,22,23^. The faster flow rates necessary for high-throughput culturing devices ^24^ could be achieved by scaling up the features of the PWM chip, which would decrease fluidic resistance.

### Characterizing the dynamic range

For all experiments we used a minimum duty cycle of 0.1 and maximum of 0.9 to ensure that the valve opening and closing times (30 ms, Supplementary Fig. 1) were not too substantial compared to pulse lengths. Cycle times greater than those tested in this work equate to longer pulsing lengths and can most likely support more extreme duty cycles. In order to broaden the dynamic range of output concentrations while keeping duty cycles between 0.1 and 0.9, we connected the chip to multiple preparations of the same substance at different concentrations. A series of 9-fold dilutions was performed to prepare 3 stock solutions of 10 kDa FITC-dextran. Our automated platform selects which stock solution to use based on the desired output concentration.

We performed this experiment using standard PWM conditions (1 s cycle length, long serpentine channel, and a 27 μL/h flow rate). The chip was programmed to create concentrations that fell within low (0.15–1.35 μM), medium (>1.35–12.15 μM), and high (>12.15–109.35 μM) ranges. The difference between the maximum (109.35 μM) and minimum (0.15 μM) concentration values is ~3 orders of magnitude, or a 729-fold change. We also acquired reference images of each stock solution, allowing us to convert fluorescence intensities at the outlet of the mixing channel to absolute concentrations. We found that the generated concentration profiles closely matched the programmed values for all 3 inlet pairs (Fig. 5). Diffusion of FITC-dextran in the serpentine channel converts the step-wise profile of the input file into a smooth contour.

**Figure 5.**
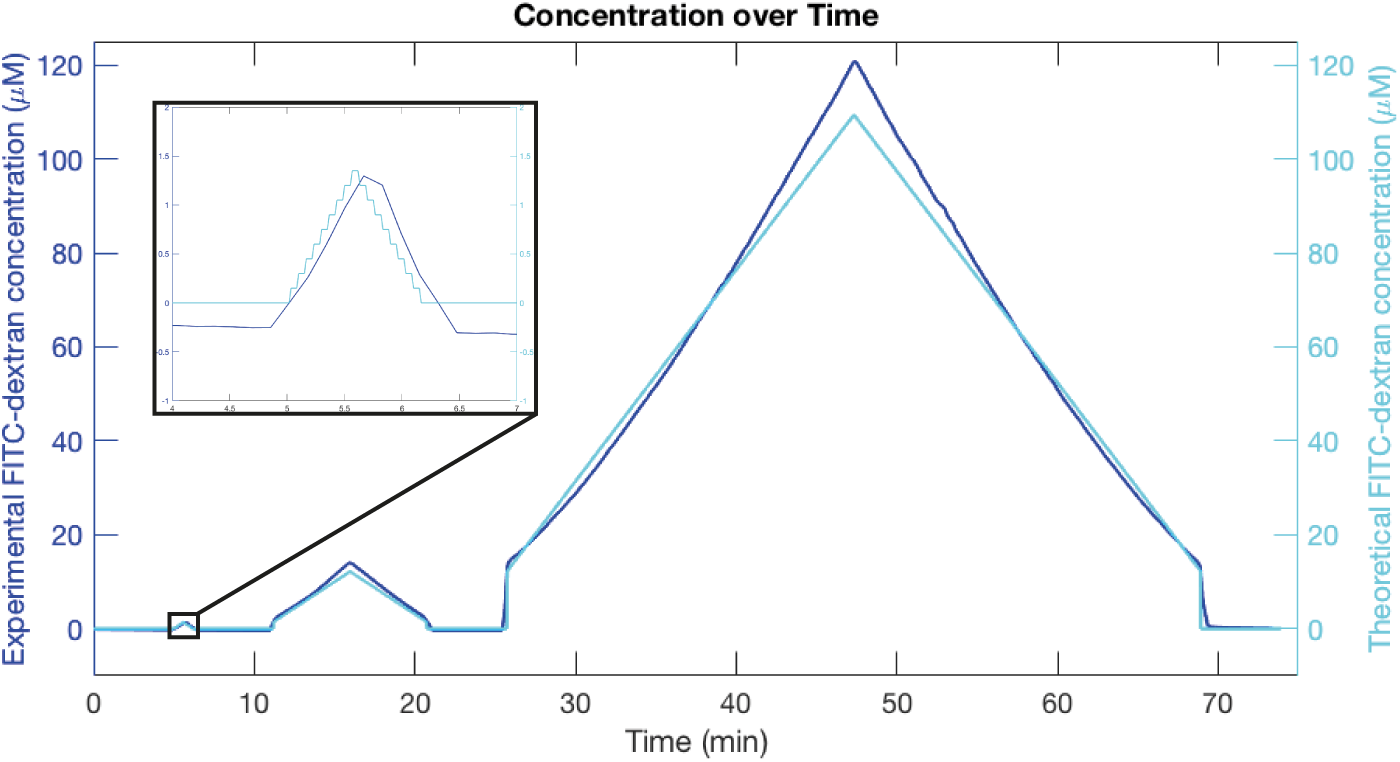
Control of on chip concentration. Experimental result superimposed with programmed values for 10 kDa FITC-dextran. Images were acquired at the end of the long serpentine channel every 10 s, analyzed for fluorescence intensity, and converted to absolute concentration using calibration images.

Due to the limited sensitivity of fluorescence detection on our setup, we tested only 3 serial dilutions. The 729-fold concentration range presented in Figure 5 is for example well within the typical difference between the minimum and maximum serum concentration of antibiotics measured in clinical studies^25,26^. However if all 6 inlets of the chip were to be utilized (5 stock solutions prepared as 9-fold serial dilutions and 1 buffer), a 59049-fold range of output concentrations could be produced. These results suggest that our technology can accurately replicate complex *in vivo* concentration profiles.

### Connecting the PWM module to a second chip

The PWM module can be easily integrated into chips fabricated by multilayer soft lithography^27^ due to its small size and the standard dimensions of its channels. To extend the use of the PWM module, we tested whether the PWM module could be connected to a second chip, bypassing the need to re-fabricate existing chips and allowing the PWM module to work with a wide variety of microfluidic devices fabricated with different techniques and materials. We constructed this setup by using flexible PEEK tubing to join the outlet of the short, straight channel (Fig. 1a) of the PWM module to the inlet of a second chip. The PEEK tubing has a 50.8 μm inner-diameter, equating to a ~1.5x larger cross-section than the microfluidic mixing channels. Faster flow rates were required to prevent excessive diffusion and delayed response times. The geometry of the PEEK tubing also leads to lower fluidic resistance compared to the microfluidic channels, allowing one to obtain higher flow rates with less pressure.

We successfully produced programmed concentration profiles in the second chip when using 10 cm of connective PEEK tubing, a medium flow velocity (47 μL/h), and a 4 s total cycle length (Fig. 6a,b). A faster flow rate of 80 μL/h was achieved when the cycle was shortened to 2 s (Fig. 6a,b). Compared to the minimum flow rate of 11 μL/h we previously demonstrated (Fig. 4), this signifies that a 7.3-fold range of flow rates can be used with our platform. The ability to support higher flow rates ensures that our PWM module can be used in conjunction with more high-throughput applications such as microfluidic chemostats^28,29,30^. The 2-chip setup can likely be used with an even greater range of flow rates so long as the total cycle time is adjusted accordingly to provide control over diffusional mixing.

**Figure 6.**
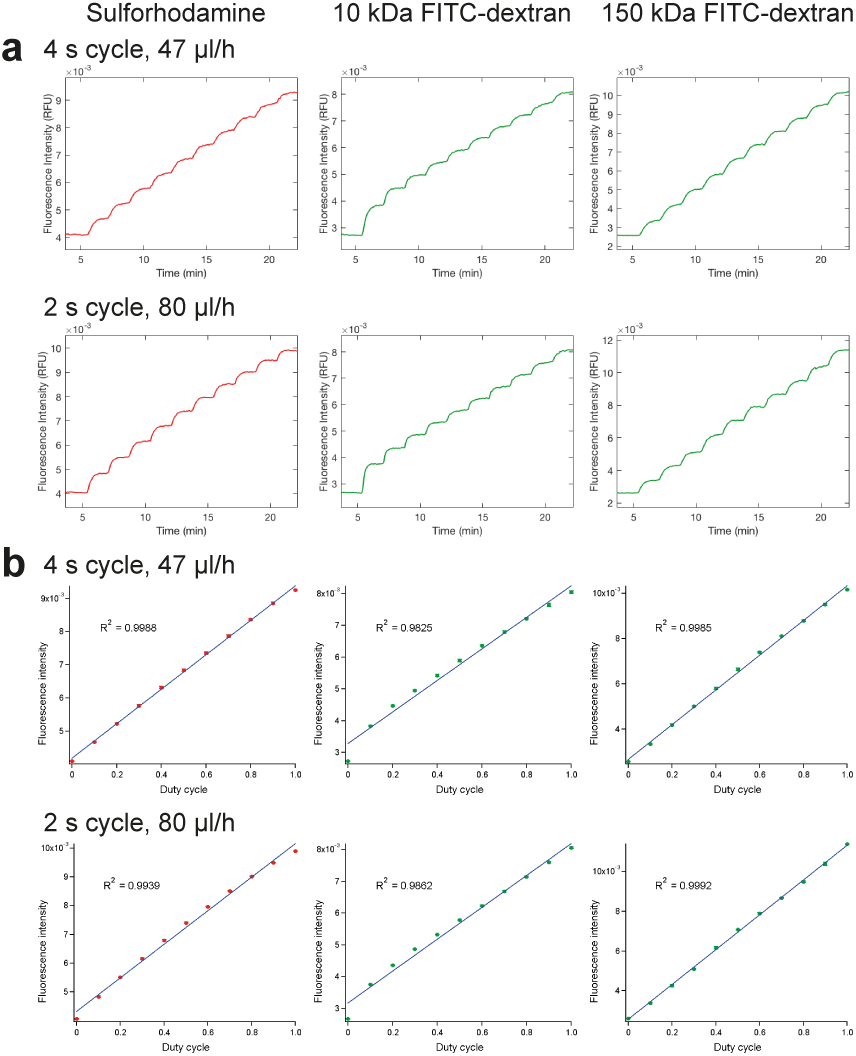
PWM chip used upstream of a second device. (a) Measured fluorescence intensity resulting from step-wise duty cycle increases over time. Images were acquired near the inlet of the second chip every 5 s. (b) Mean fluorescence intensity as a function of programmed duty cycle. Error bars represent standard deviations. Means were calculated from 10-15 RFU measurements and did not include the data points that fell between two concentrations (duringramping up).

### Complex, long-term and PK/PD concentration profiles

Complex concentration profiles can also be implemented when a PWM module is connected upstream of a second chip. Using a flow rate of 47 μL/h, a 4 s cycling time, and 6 cm of PEEK tubing to connect the two chips, we simultaneously changed the concentrations of two substances, generating sine curves of changing periods for sulforhodamine and 10 kDa FITC-dextran (Fig. 7a). The 4 s cycle consisted of 3 components: a sulforhodamine pulse, a FITC pulse, and a buffer pulse (Fig. 7a). Sulforhodamine and FITC could each be pulsed for a maximum of 2 s, and pulse on/off times were selected independently for each substance. The off times for both sulforhodamine and FITC were combined into one buffer pulse. This technique allowed us to create sine curves with periods of 5, 10, 20, 30, and 45 min for FITC, and 80 and 60 min for sulforhodamine (Fig. 7a). Our LabVIEW interface enables the user to run custom concentration profiles encoded in text files and is not limited to simple mathematical functions. The ability to independently control the concentration of several substances at once on chip is pertinent to complex biological studies that require the simultaneous delivery of multiple factors as commonly required in stem cell differentiation or reprogramming experiments^31^.

**Figure 7.**
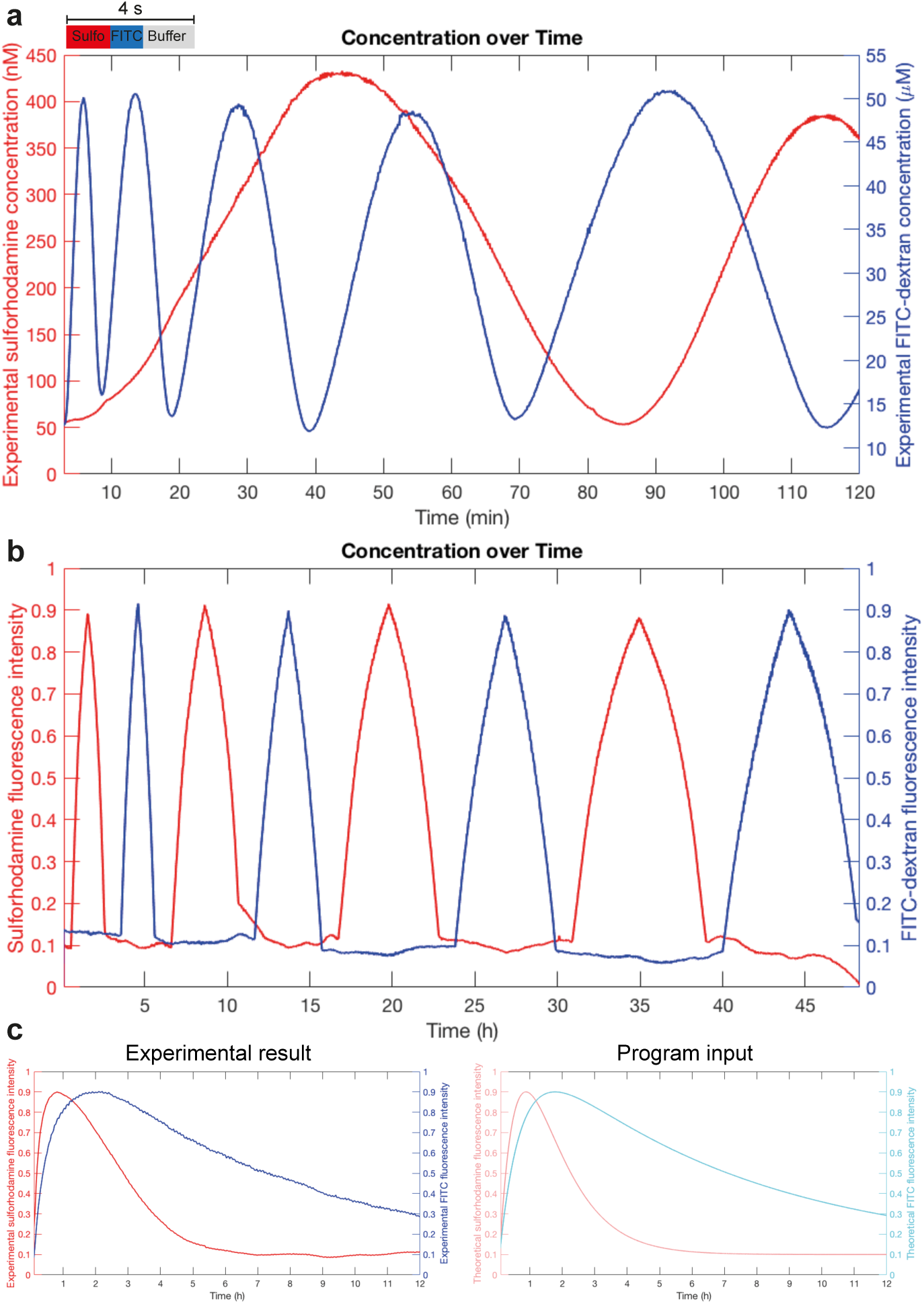
Complex experiments on the PWM chip. (a) PWM of sulforhodamine and 10 kDa FITC-dextran in parallel, programmed as sine curve functions with changing periods. Images were acquired near the inlet of the second chip every 10 s, analyzed for fluorescence intensity, and converted to absolute concentration using calibration images. (b) PWM of sulforhodamine and 10 kDa FITC-dextran in parallel, gradually ramped to create peaks of 2,4,6, or 8 h duration. Images were acquired at the end of the long serpentine channel every 2 min for 48 h, analyzed for fluorescence intensity, and normalized from 0 to 1 based on minimum and maximum fluorescence intensity. (c) Simultaneous generation of two different PK profiles using sulforhodamine and 10 kDa FITC-dextran. Images were acquired and analyzed as in (b).

We have shown that our device can implement fast concentration changes and programs lasting up to 2 h (Fig. 3, Fig. 7a). These parameters are compatible with chemical and *in vitro* studies. However, biological studies and especially those involving cell cultures call for extended experimental periods. We thus evaluated the capacity of our device to manipulate and sustain concentrations on time scales spanning up to 48 hours. Using the long serpentine channel, a flow rate of ~30 μL/h, and a 4 s cycling time, we performed PWM with sulforhodamine and 10 kDa FITC-dextran in parallel. Each substance was either stably maintained at 0.1 (10% of maximum) or gradually ramped up to, or down from, 0.9x (90% of maximum) (Fig. 7b). Each peak spanned 2, 4, 6, or 8 h, which are time ranges relevant for drug dosage and pharmacokinetic studies^25,26^. We found that for all ramping times, the desired profile was accurately generated (Fig. 7b). The 48 h experiment was not subject to errors or chip failure, showing the robustness of our setup.

The ability to slowly ramp concentrations over long periods of time would be relevant for on chip pharmacokinetic and pharmacodynamics (PK/PD) studies. To demonstrate the utility of our device for this application, we mimicked the 12 h concentration time-profiles in plasma for orally administered drugs^32^ (Fig. 7c). Chip operating conditions were the same as those employed in Fig. 7b, and sulforhodamine and FITC were programmed simultaneously. Both molecules started at 0 concentration and quickly increased to 0.9 (90% intensity), corresponding to release of the drug from the formulation and absorption into the bloodstream^33,34^. After reaching maximum concentration, both substances decrease gradually (to a final value of 0.1 for sulforhodamine and 0.29 for FITC), representing metabolism and elimination of the drug. The experimental results closely match the programmed values in both shape and concentration (Fig. 7c). These profiles can be custom-generated on chip to reflect different drugs, delivery methods, and formulations. This feature would be especially advantageous for studies investigating the interactions of multiple drugs to identify synergy or antagonism. When used in this manner, our PWM module could be connected upstream of existing devices to facilitate PK/PD and other dosage-dependent studies. Compared to previously developed PWM microfluidic chips^9,10,11^, our module enables more complex studies to be performed (Supplementary Table 1). We have shown that our chip can be used in a variety of configurations, and a guide to selecting the optimal conditions for each desired flow rate and experiment type is presented in Figure 8.

**Figure 8.**
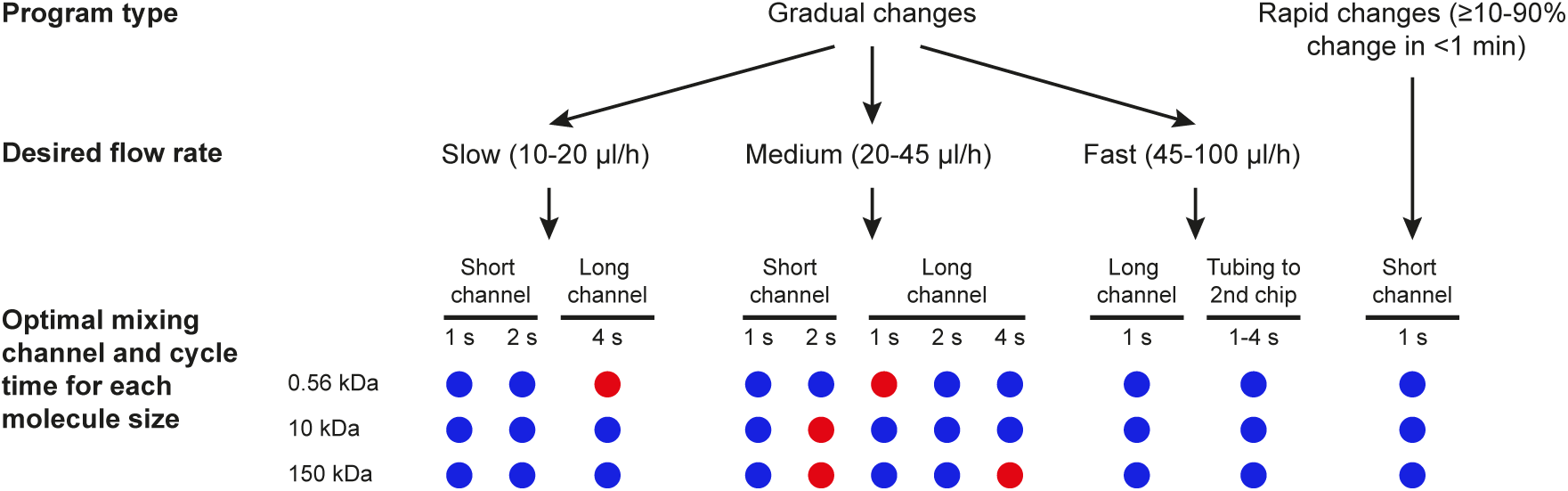
Guide to selecting chip operating conditions for different experimental setups. Blue, optimal; red, non-optimal.

## Discussion

Our PWM module demonstrates several improvements over current microfluidic dilution methods (Supplementary Table 1). Compared to previous chips, our platform enables long-term experiments, manipulates the concentration of multiple substances in parallel, and works with a variety of molecule sizes and flow rates. Moreover, the PWM module can easily be connected upstream of a second device. We have thoroughly characterized the platform to aid the user in selecting the optimal conditions for a given experiment (Fig. 8). The PWM module itself is easy to fabricate using standard multilayer soft lithography and the design files are available for download (lbnc.epfl.ch).

Microfluidics is increasingly being used as a tool to model *in vivo* conditions due to reduced sample requirements, regulation of the microenvironment, and compatibility with single-cell analysis. However, typical devices lack the ability to gradually ramp concentrations on chip and instead simply perform non-physiological step-functions by switching between on and off states for a given substance in the culturing medium. We address this limitation by allowing the user to generate arbitrary concentration-time profiles of several substances with complete control over both timing and concentration. The device can be programmed to generate PK/PD profiles, a feature that is especially relevant in light of recent developments in organ-on-chip and microfluidic stem cell technologies^35,36^. Overall, our PWM platform complements existing biomedical microfluidic devices and will facilitate studies that better reflect the *in vivo* environment by emulating physiologically relevant temporal changes in concentrations.

## Acknowledgments

This work was funded by EPFL. We thank Nadanai Laohakunakorn and Mathieu Quinodoz for help with LabVIEW programming. We thank Gauthier Croizat for helpful discussions and assistance with fluidic modeling.

## Experimental

### Chemicals

Sulforhodamine, FITC-dextran 10 kDa, and FITC-CM-dextran 150 kDa were purchased from Sigma Aldrich.

### Automated pneumatic setup

Actuation of microfluidic valves was controlled by a setup consisting of a 16 channel USB relay module (Numato lab) connected to 24V solenoid pneumatic valves arranged on manifold (Pneumadyne). A custom written LabVIEWVI was used to control the opening and closing of the valves on the chip. To run the program, the user supplies a text file containing a list of desired concentrations over time. The LabVIEW program selects which flow inputs to use, then calculates and executes the pulsing times of each input required to achieve the desired concentration. The total cycle length can be adjusted in the program.

### Imaging

Imaging of the microfluidic chip was performed on a Nikon Ti-E Eclipse automated microscope using NIS Elements. Images were acquired with an Ixon DU-888 camera (Andor Technology), using 20x magnification to capture the last ~700 μm of the mixing channel in the field of view. For imaging fluorescence, two HC filter cubes were used: TexasRed (HC 562/40, HC 624/40, BS 593) for sulforhodamine, and FITC (HC 482/35, HC 536/40, BS 506) for FITC-dextran (all filters from AHF Analysentechnik AG). Images were analyzed using a custom written Matlab script in which a box of 100 μm length and including the entire width of the channel was selected and analyzed for mean intensity. High-speed imaging of valve closure was performed using a FasTec Imaging HiSpec 2G camera and FasTec imaging software.

### Microfluidic device fabrication

Microfluidic chips were fabricated as previously described^37^. Devices were designed in Clewin (WieWeb software, Netherlands). Two molds were designed: one for the control layer, which contains the valves, and another for the flow layer, which contains the channels and chambers necessary for reagent introduction and cell culturing. The control layer was scaled by 101.5% to account for PDMS shrinkage during curing. The molds were fabricated using standard photolithography methods. The control layer mold was patterned with SU-8 photoresist (Gersteltec, Switzerland) to a height of 30 μm, then exposed and developed. The channels on the flow layer were generated by spin coating AZ9260 to a height of 14μm, followed by exposure and development. The AZ9260 was annealed at 120°C for 10 min to produce the rounded profile that is required for complete valve closure. Polydimethylsiloxane (PDMS;Sylgard 184, Dow Corning Corp., USA) was cast onto the molds and multilayer soft lithography techniques were used to assemble the chip. A thick layer of PDMS (5:1 ratio of parts A:B) was poured onto the control layer, whereas the flow layer was spin coated with PDMS (20:1 ratio of parts A:B) with a ramp of 15 s and a spin of 35 s at 3000 rpm. The molds were baked for 30 min at 80°C. The control layer chips were then cut, removed from the mold, and punched with inlet holes. The control layer chips were aligned to the flow layer, and the assembly was baked for 90 min at 80° C. The aligned devices were then cut, removed from the mold, and punched with inlet and outlet holes. The chips were placed on top of a glass slide that had been coated with 20:1 PDMS (15s ramp time and 35s spin at 1500 rpm) and baked for 30 min at 80°C. Finally, the entire assembly was placed under a weight and bonded overnight at 80°C.

### Microfluidic chip operation

The pressures of the flow and control layers of the chip were controlled by using a custom built pneumatic setup. The valves of the control layer were first primed with filtered water at 5 psi. Once all air had been removed from the control lines, the control layer pressure was increased to 25 psi. The flow layer of the chip was washed with water before each experiment. Flow rate was determined by measuring the volume of liquid exiting the chip over a period of time. Tygon tubing was used for most fluidic connections. For experiments in which a PWM device was attached to a second device, 50.8 μm diameter PEEK tubing (VWR) was used. One end of a 6-10 cm piece of the tubing was inserted directly into the outlet hole of the PWM chip and the other end was placed into the inlet of the second chip.

